# Intronic CNVs cause gene expression variation in human populations

**DOI:** 10.1101/171165

**Authors:** Maria Rigau, David Juan, Alfonso Valencia, Daniel Rico

## Abstract

Introns can be extraordinarily large and they account for the majority of the DNA sequence in human genes. However, little is known about their population patterns of structural variation and their functional implication. By combining the most extensive maps of CNVs in human populations, we have found that intronic losses are the most frequent copy number variants (CNVs) in protein-coding genes in human, with more than 12,986 intronic deletions, affecting 4,147 genes (including 1,154 essential genes and 1,638 disease-related genes). These intronic length variation results in dozens of genes showing extreme population variability in size, with 40 genes with 10 or more different sizes and up to 150 allelic sizes. Intronic losses are frequent in evolutionarily ancient genes that are highly conserved at the protein sequence level. This result contrasts with losses overlapping exons, which are observed less often than expected by chance and almost exclusively affect primate-specific genes. An integrated analysis of CNVs and RNA-seq data showed that intronic loss can be associated with significant differences in gene expression levels in the population (CNV-eQTLs). These intronic CNV-eQTLs regions are enriched for intronic enhancers and can be associated with expression differences of other genes showing long distance intron-promoter 3D interactions. Our data suggests that intronic structural variation of protein-coding genes can exert an important role in maintaining the variability of gene expression and splicing in human populations.

**Author Summary:** Most human genes have introns that have to be removed after a gene is transcribed from DNA to RNA because they not encode information to translate RNA into proteins. As mutations in introns do not affect protein sequences, they are usually ignored when looking for normal or pathogenic genomic variation. However, introns comprise about half of the human non-coding genome and they can have important regulatory roles. We show that deletions of intronic regions appear more frequent than previously expected in the healthy population, with a significant proportion of genes with evolutionary ancient and essential functions carrying them. This finding was very surprising, as ancient genes tend to have high conservation of their coding sequence. However, we show that deletions of their non-coding intronic sequence can produce considerable changes in their locus length. We found that a significant number of these intronic deletions are associated with under- or over-expression of the affected genes or distant genes interacting in 3D. Our data suggests that the frequent gene length variation in protein-coding genes resulting from intronic CNVs might influence their regulation in different individuals.

## Introduction

Most eukaryotic protein coding genes contain introns that are removed from the messenger RNA during the process of splicing. In humans, up to 35% of the sequenced genome corresponds to intronic sequence, while exons cover around the 2.8% of the genome (based on the genome version and gene set used for this study). Human introns can have very different lengths, contrarily to exons. This difference in intron length leads to substantial differences in size among human genes, which cause differences in the time taken to transcribe a gene from seconds to over 24 hours [1]. Indeed, intron size is highly conserved in genes associated with developmental patterning [2], suggesting that genes that require a precise time coordination of their transcription are reliant on a consistent transcript length. It has been suggested that selection could be acting to reduce the costs of transcription by keeping short introns in highly expressed genes [3], which are enriched in housekeeping essential functions [4]. Genes transcribed early in development [5–7] and genes involved in rapid biological responses [8] also conserve intron-poor structures. Interestingly, Keane and Seoighe [9] recently found that intron lengths of some genes tend to coevolve (their relative sizes co-vary across species) possibly because a precise temporal regulation of the expression of these genes is required. In fact, these genes tend to be coexpressed or participating in the same protein complexe [9].

It is well known that introns contribute to the control of gene expression by their inclusion of regulatory regions and non-coding functional RNA genes or directly by their length [10–12]. Despite the importance of introns in regulating transcription levels, transcription timing and splicing, little attention has been payed to their potential role in human population variability studies. A recent analysis of the literature has revealed a substantial amount of pathogenic variants located “deep” within introns (more than 100 bp from exon-intron boundaries) which suggests that the sequence analysis of full introns may help to identify causal mutations for many undiagnosed clinical cases [13]. Given that direct associations between intronic mutations and certain diseases have been reported [13–16], we need to characterise the normal genetic variability in introns so we can better distinguish normal from pathogenic variations.

## Results

### Deletions are enriched in purely intronic regions

We studied the effect of structural intronic variants on protein coding gene loci in healthy humans using five copy number variant (CNV) maps of high resolution [17–21]. Most of these CNVs were detected using whole genome sequencing (WGS) data, which allows to determine the exact genomic boundaries of these variants. CNVs may have neutral, advantageous or deleterious consequences [22] and can be classified in ***gains*** (regions that are found duplicated when compared with expected number from the reference genome, which is 2 for autosomes), ***losses*** (homozygously or heterozygously deleted regions) and ***gain/loss CNVs*** (regions that are found duplicated in some individuals - or alleles - and deleted in others). Each of the maps in our study was derived from a different number of individuals, from different populations and using different techniques and algorithms for CNV detection (**Fig S1** and **Table S1)**. Due to these differences, each dataset provided us with a different set of CNVs (**Fig S1**), which we analysed independently, excluding sex chromosomes and private variants.

CNVs affect genes in different ways depending on the degree of overlap with them. Some CNVs cover entire genes (from now on ***whole gene CNVs***), other CNVs overlap with part of the coding sequence but not the whole gene (***exonic CNVs***) and other CNVs are found within purely intronic regions (***intronic CNVs***, not overlapping with any exon from any annotated isoform, **Fig 1A**). The latter group is the most common, with 63% of all CNVs falling within intronic regions, but remains the least studied. More than the 95% of these 12,986 intronic CNVs are losses (12,334) or gain/loss CNVs (652). The prevalence of losses in introns is in stark contrast with whole gene CNVs (1,412), which tend to be exclusively gains (55% of the cases) or gain/loss CNVs (25% of the cases) (**Fig 1B-D**).

**Fig 1.**
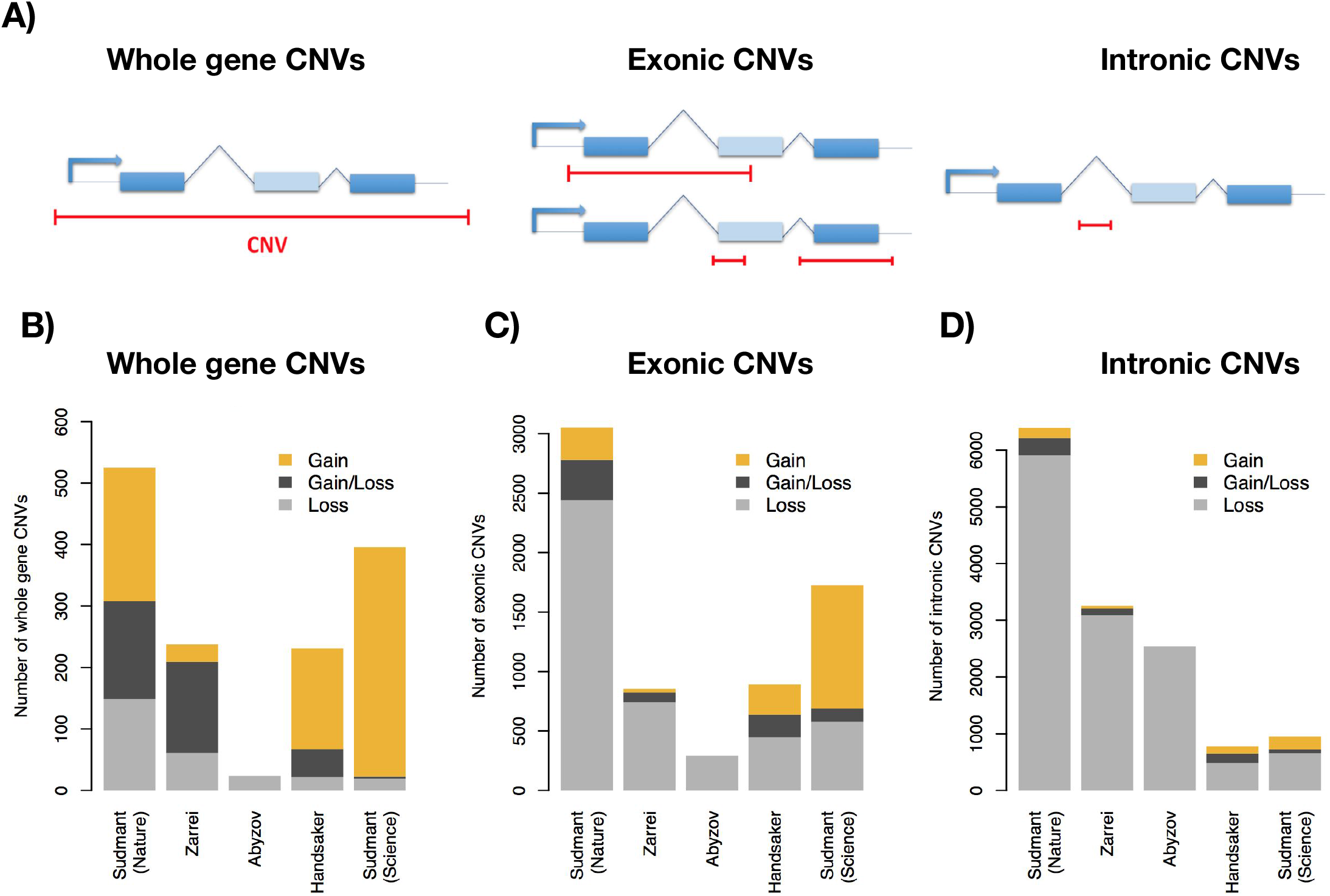
Types of CNVs in the different datasets. (A) CNVs can overlap entire genes or fractions of genes. CNVs overlapping with exons of a gene (exonic CNVs) and CNVs found within introns (intronic CNVs). (B-D) Number of whole gene, exonic and intronic CNV events, showing the different proportions of CNV gains, losses and gain and loss CNVs.

Surprisingly, purely intronic losses are not only the most prevalent form of CNV, but also they are observed more often than expected by chance (**Table S2**). We compared the observed values with expected distributions calculated using permutations in local and global background models (see Methods and **Fig S2**). We find significantly more deletions (4.14-9.3%) falling in introns than expected in 3 out of the 5 maps (4.14% in Sudmant-Nature, *P* = 0.0002, global permutation test). For the sake of clarity, the P-values in the main text correspond to the results in Sudmant-Nature’s map [20] using the global background model (unless otherwise indicated). The results obtained with the alternative background model and with both models in additional maps are shown in the supplementary tables and figures.

In contrast with intronic deletions, there are 51.2% fewer coding deletions (overlapping with exons) than would be expected by chance (*P* < 1e-04, Sudmant Nature, global permutation test). These patterns are consistent using the two different background models (**Fig S2** and Table **S2**) and are independent of intron size **(Fig S3)**.

The enrichment of deletions in introns might seem contradictory to what was originally reported by the 1000 Genomes (1KG) Project [20], as they stated that introns had *less* CNVs than expected by chance. However, we would like to note that they did not separate purely intronic from intron-exon overlapping deletions, while we are talking about strictly intronic deletions (see Methods for details). Indeed, if we group all purely intronic and intron-exon overlapping deletions together, we also observe a significant depletion (**Table S2, Fig S2**).

### Highly variable sizes in highly conserved protein-coding genes

The percentage of each intron that can be lost in the population due to CNV losses is highly variable, from 0.03% to 96.8% (51 bp to 293 kb), representing a loss of the 0.01% to 77.5% of the total genic size (51 bp to 893,4 kb, **Fig 2A-C**). Some examples of genes with a notable change in size after a single intronic deletion in one individual are the neuronal glutamate transporter *SLC1A1 (Solute Carrier Family 1 Member 1)*, with a loss of the 37% of its genic size (**Fig 2D**) and the *LINGO2* (*Leucine Rich Repeat And Ig Domain Containing 2*, alias *LERN3* or *LRRN6C*) gene with a loss of the 34% of its size.

**Fig 2.**
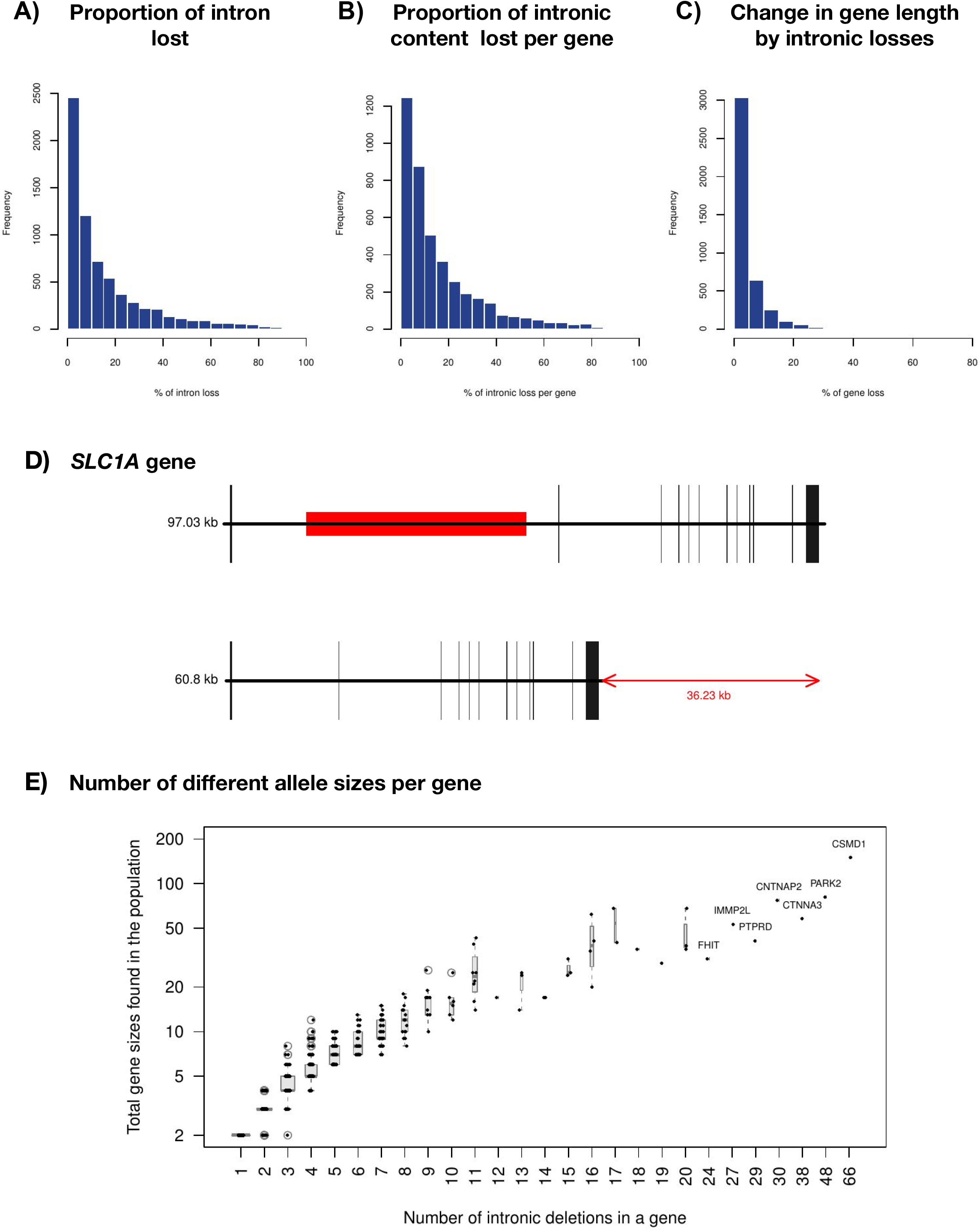
Changes in intron and gene size. (A) Proportion of the reference intron that has been observed as deleted in any of the studies. (B) Proportion of the whole intronic content of a gene that has been observed as deleted. (C) Change in gene length by intronic deletions. (D) Example of gene with a substantial change in gene size with a single intronic deletion. (E) Number of different gene sizes observed in the population as a function on the number of intronic deletions detected. Genes names of the seven most extreme cases are indicated.

The combination of different intronic deletions within a gene can give place to alleles of several different sizes (**Fig 2E**). Following with the same two examples, in the dataset from the final phase of the 1KG Project [20], we found 5 different intronic deletions in *SLC1A1*. These deletions result in 8 different sizes of genes in the population, with individual losses ranging from 1,1kb bp to 48kb. In *LINGO2*, the 20 different deletions give place to 36 different gene lengths in the 1KG population, with losses of 51 to 233 kp. The gene with more different allele sizes in the 1KG population [20] is *CSMD1* (*CUB And Sushi Multiple Domains 1*), with a total of 66 common intronic annotated deletions that, combined, produce 150 alleles of different sizes. Strikingly, *CSMD1* is highly conserved at the protein level and is amongst the most intolerant genes to functional variation. According to the ranking of the RVIS (Residual Variation Intolerance Score) gene scores [23], which is based on the amount of genetic variation of each gene at an exome level, only 0.169% genes in the human genome are more intolerant to variation in their coding sequence than *CSMD1*. In summary, intolerance to variation in the coding sequence seems to be compatible with extreme variation in the intronic sequence. These losses might affect their regulation without affecting their protein structure.

A total of 1,638 OMIM genes carry intronic deletions in the population. Diseases associated to *SLC1A1* (OMIM: 133550) include Dicarboxylic Aminoaciduria and susceptibility to Schizophrenia, while *LINGO2* (OMIM: 609793) has variants associated with essential tremor and Parkinson disease and also has an intronic SNP associated with body mass [24]. *CSMD1* has been associated to diseases such as Benign Adult Familial Myoclonic Epilepsy (Malacards [25], MCID: BNG079) and Smallpox (MCID: SML019). Interestingly, rare intronic deletions in this gene have been recently reported to be associated to both male and female infertility [26]. To better understand possible epistatic effects between protein-coding and intronic mutations, it will be useful to incorporate information about gene length variation in future studies of these disease genes.

### Intronic deletions are frequent in evolutionary ancient and essential genes

Structural variants in the germline DNA constitute an important source of genetic variability that serves as the substrate for evolution. Therefore, dating the evolutionary age of genes allows the study of structural variants that were fixed millions of years ago. Whole gene CNVs are known to differentially affect genes depending of their evolutionary age, mainly involving evolutionary young genes [27]. Genes of younger ages are generally cell-type specific, while ancient genes tend to be more conserved, ubiquitously expressed and enriched in cellular essential functions. Intrigued to see many cases where intronic CNVs were affecting highly conserved protein-coding loci, we compared the distribution of coding (including exonic and whole gene) and intronic deletions across different gene ages (**Fig 3**). These and subsequent analyses were done using 3 maps: Sudmant-Nature’s [20], Zarrei’s [19] and Abyzov’s [17] maps. Handsaker’s [18] and Sudmant-Science’s [21] maps were discarded because they had very few intronic deletions (less than 1000, **Fig S1**).

**Fig 3.**
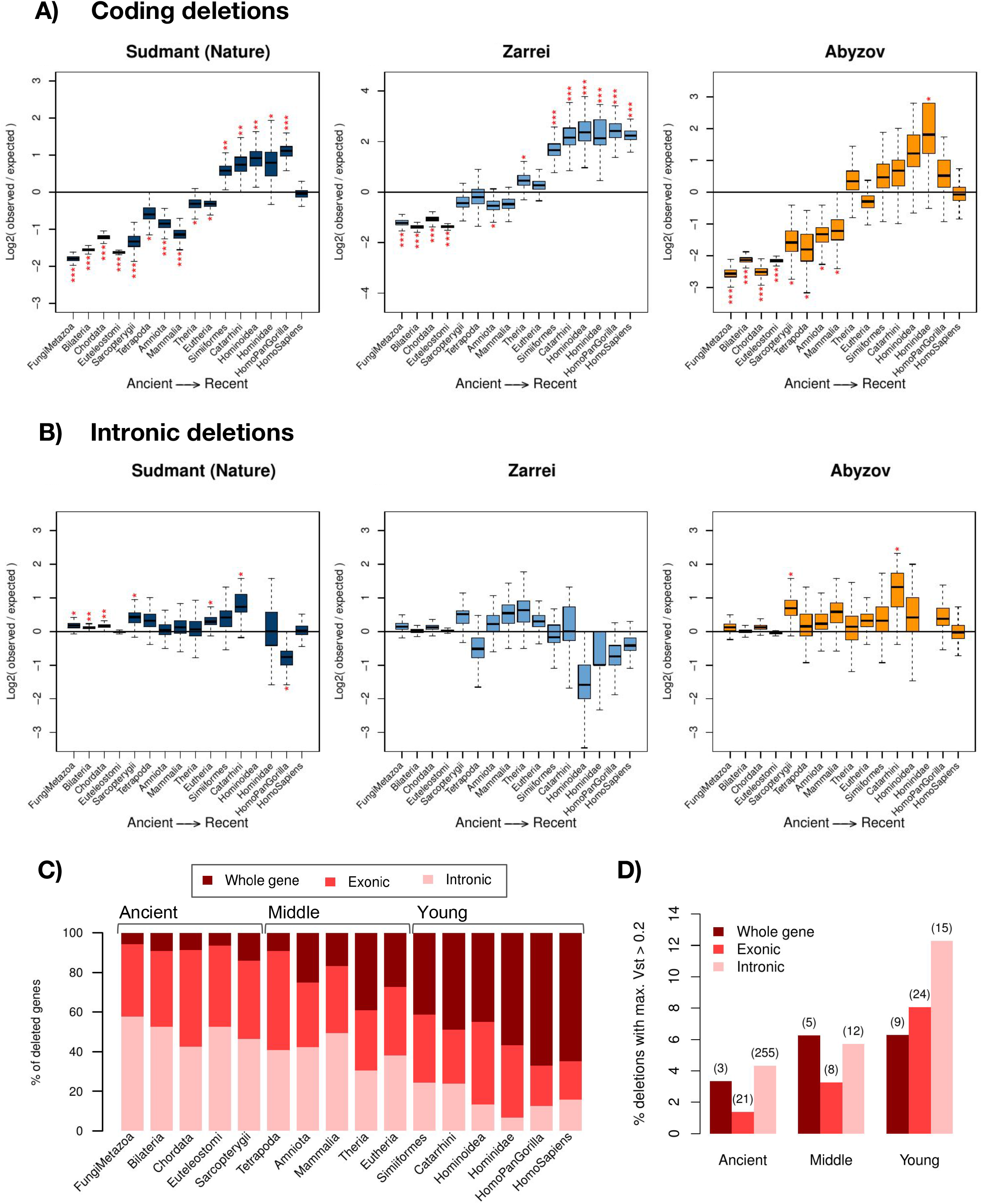
Evolutionary age of affected genes. Ratios of observed versus expected number of genes from each gene evolutionary age that contain deletions overlapping with exons, including partial and whole gene CNVs (A) or intronic deletions (B). Expected values were calculated with 10,000 random permutations using a global background model. Red asterisks mark the significantly enriched groups of genes. Significance: * for P<0.05, ** for P<0.005 and *** for P<0.0005. Plot (C) shows, from all the genes overlapping with deletions after aggregating the three maps, what is the proportion of genes that have all or part of their exons affected by deletions and what is the percentage of genes with intronic deletions only. Bar width is proportional to the percentage of genes from each evolutionary age that is affected by deletions of any kind, which spans from 18.5% (Mammalia) to 49.8% (HomoPanGorilla). The equivalent figure for each separate map is shown in Fig **S7**. **(D)** Percentage of highly stratified variants (HSV, maximum Vst > 0.2) in each age group and by type of overlap with the gene. The absolute number of deletions is indicating above each bar.

We observed that most ancient genes are depleted of deletions that affect their coding regions, while primate-specific genes are enriched with coding CNVs (**Fig 3A**). This pattern was also observed when CNV gains were included (**Fig S4**), meaning that the coding region of recent genes have a higher tendency to be lost or disrupted. The generation of random background models revealed that ancient genes (present in the *Sarcopterygii* ancestor) were significantly depleted of coding region losses (both exonic and whole gene, *P* < 1e-04, global permutation test), while these were enriched in young genes (from *Hominoidea* to *Homo sapiens*, *P* < 1e-04, global permutation test; see **Fig 3A** and **Table S2**).

In contrast with coding deletions, we found that the proportion of ancient genes with intronic deletions was higher than that of young genes (**Fig S5**), and also higher than expected by chance in Sudmant-Nature’s map (*P* = 2e-04, global permutation test, **Fig 3B, Table S2**). This finding was confirmed with additional analyses considering only genes with big introns (larger than 1,500 bp, see **Table S2**) and by calculating the enrichment within big introns independently from genes (**Fig S6 and Table S2)**. In summary, the frequency of coding deletions in ancient genes is lower, but their frequency of intronic deletions is higher than the one observed for young genes (**Fig 3C and Fig S7**).

We would expect that essential genes, which tend to be ancient [28], could be an exception to the enrichment of deletions. Essential genes have on average shorter introns than the rest of the genes [29,30] and relative to the genes of the same evolutionary age (**Fig S8A** and **B**). Up to 1,154 essential genes carry intronic deletions if we take into account all five CNV maps. In Sudmant Nature, 907 essential genes have intronic deletions, a higher number than expected by chance (*P* = 0.034, global permutation test, **Table S2**).

We investigated if intron variability in genes was associated with any biological function. Genes with more or less intronic deletions than expected by chance (**Table S3**, see Methods) were not associated to any particular function using DAVID [31]. Nevertheless, genes with less intronic deletions than expected show more protein-protein interactions among them than expected by chance (P = 2.43e-10, calculated with STRING [32]). These results are compatible with previous evolutionary studies that showed high levels of conservation of intron length in genes associated with development protein complexes in mammals [2], presumably to facilitate a more precise temporal regulation of expression [9].

Population stratification of CNVs has previously been suggested to be indicative of loci under adaptive selection [20,21]. We identified 352 highly stratified variants (HSVs, maximum Vst>0.2, see Methods) from Sudmant-Nature’s map overlapping with protein-coding genes: 282 are intronic, 53 exonic and 17 whole gene. We classified deleted regions according to the age of the genes and the type of gene structure affected and calculated the percentage of each group that is highly stratified (**Fig 3D**). Interestingly, the contribution of intronic HSVs is higher for younger genes, a pattern coherent with the expected higher functional impact of HSVs in older genes. Remarkably, the percentage of intronic HSVs is similar or higher than that of whole-gene and exonic HSVs in all age groups (and always higher than partial exonic deletions). These signatures of positive selection in purely-intronic CNVs suggest that a fraction of them might contribute to human adaptation.

### Intronic deletions are associated with gene expression variability in the population

Multiallelic CNVs affecting whole genes have been shown to correlate with gene expression: generally, the higher the number of copies of a gene, the higher its expression levels [18,20]. We hypothesized that intronic size variation may also impact the expression of the affected genes (without affecting the actual number of copies of the gene). Therefore, we looked into the possible effect of intronic hemizygous deletions on gene expression variation at the population level, comparing the effects with hemizygous deletions in coding (whole gene and exonic) and intergenic non-coding deletions (**Fig 4**). We used available RNA-seq data from Geuvadis [33] that was derived from lymphoblastoid cell lines for 445 individuals for whom we have the matching CNV data from the 1KG Project [20].

**Fig 4.**
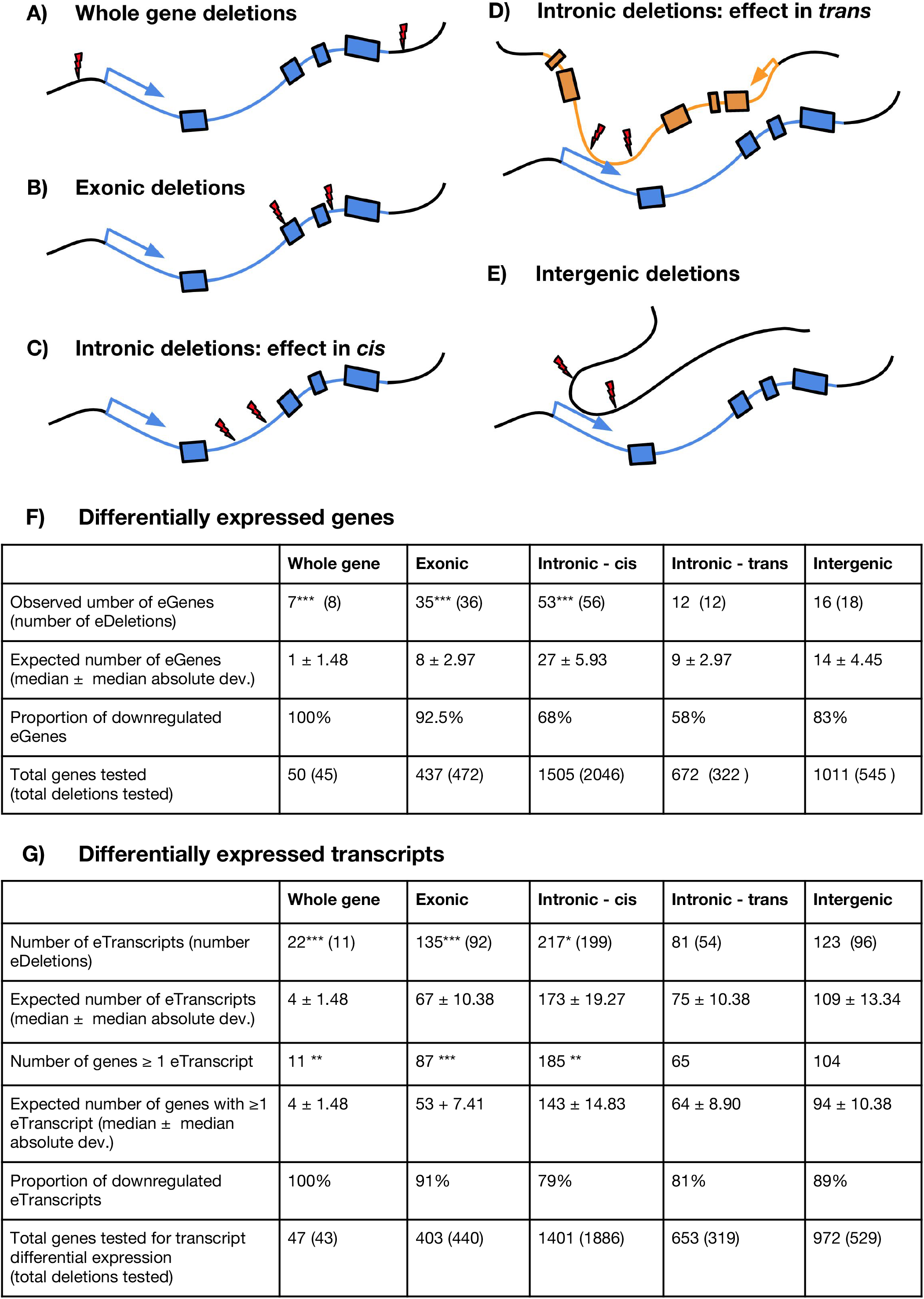
DEL-eQTLs. (A-E) Five types of DEL-eQTLs analysed. Thunder symbols indicate deletion breakpoints. (F) DEL-eQTL results for the five types. Number of eGenes, eTranscripts and genes with eTranscripts when comparing expression levels of individuals with a reference allele and an allele with a specific deletion versus individuals with two reference alleles. P-values obtained after performing Student’s t-tests were FDR-corrected (FDR = 10%). The number of expected eGenes, eTranscripts or genes with eTranscripts was calculated after randomizations of the individuals carrying or not the deletion, and P-values were calculated by comparing the observed versus the 10000 random values. Significance: * for P<0.05, ** for P<0.005 and *** for P<0.0005. (G) Examples of intronic deletions in a gene associated to expression changes of another gene that interacts in 3D. Black boxes represent exons and the light blue box the PCHiC fragment in contact with the differentially expressed gene. The position of the deletion is marked with dashed red lines. Gene expression of the eGenes is represented using PEER-RPKM values.

In order to look for differences in gene expression we selected variants for which we had at least 2 hemizygous individuals (individuals with copy number = 1) and at least 2 wild-type individuals (copy number = 2) and we compared the expression levels among these two groups to identify deleted CNV regions associated with expression quantitative trait loci (eQTL, **Fig 4F**). We will refer to the deleted regions associated with expression changes as DEL-eQTLs, and the genes associated with as eGenes. For comparative purposes, we first looked at the effect of hemizygous deletions in coding regions (whole gene and exonic DEL-eQTLs). We found that 7 eGenes out of 50 genes with whole gene deletion CNVs resulted in significant downregulation of gene expression in lymphoblastoid cell lines (14%, a higher number eGenes than expected by chance, *P* < 5e-4, permutation test, **Fig 4F**). In addition, we found 35 eGenes out of 437 genes with partial exonic deletions that were differentially expressed (8%, a number higher than expected by chance, *P* < 5e-4, permutation test, **Fig 4F**). The majority of these eGenes (32/35) where down-regulated in the individuals carrying the deletion.

Although intronic deletions do not affect the coding sequence of genes, we observed significant differences in gene expression in 53 eGenes out of the 1,505 genes with intronic deletions, a number of intronic-eGenes that is also higher than expected by chance (*P* < 51e-4, permutation test) **(Fig 4F**). Given the higher abundance of intronic deletions in the population, the absolute number of intronic-eGenes (53 genes) was similar to the total of coding-eGenes (39 genes, **Fig 4F** and **Table S4**). Of the intronic-eGenes, 62% were downregulated and the other 38% upregulated, suggesting that intronic deletions might result both in enhancing or repressing gene expression (while coding losses mostly associate to gene down-regulation). Regulatory regions are known to be preferentially located in first introns [34]. From all 56 intronic eDeletions that are associated to changes of gene expression in our study, 17 (30.4%) are found within first introns. However, this percentage is not significantly higher than in non eDeletions (26%, P = 0.54, Fisher’s test). Finally, we identified that four of the intronic cis-eDeletions in lymphoblastoid cells are HSVs, suggesting adaptive potential of these expression differences. These intronic HSVs are located in four ancient genes (Sarcopterygii or older): *EXOC2*, *SKAP2*, *PTGR1* and *PHYHD1* (**Fig S9**). *EXOC2* is an essential gene encoding one of the proteins of the exocyst complex and is among the top 5% most conserved genes in human (RVIS = 3.34).

Since intron length can impact the inclusion of alternative exons [35], we hypothesised that there might be genes with differentially expressed transcripts (eTranscripts) in any gene containing an intronic deletion. In addition to the 53 intronic-eGenes, we found 217 introniceTranscripts in a total of 185 genes (this is more than expected by chance, *P* = 0.018, permutation test, **Fig 4F** and **Table S5)**. These results suggest that deletions within introns may cause the inclusion or exclusion of exons and thus influencing the relative proportion of alternative transcripts in many genes.

Changes in GC content as the result of intronic deletions might also contribute to these splicing differences, as in genes with long introns, the recognition of introns and exons by splicing machinery is based on their differential GC content [36,37] and the lower GC content in introns facilitates their recognition. We found that, in general, the deleted sequences have a significantly higher GC content to that of the introns where they are located (P = 1.8e-28, paired Student’s t-test), and the loss of these sequences causes a significant decrease of the overall GC content of the introns (*P* = 2.23e-16, paired Student’s t-test) (**Fig S10** and **Fig S11**). This drop of GC content is more pronounced in introns with deletions originated through transposable element insertion (TEI, P = 2.01e-9, paired Student’s t-test). The 84% of TEI deletions overlap almost completely with Alu elements (Fig S12), which are known to be GC rich. The GC drops happening in introns with deletions associated to nonallelic homologous repair (NAHR) are less significant (P = 0.0063), while the difference is not significant in deletions caused by non-homologous repair (NH) (P = 0.7676). The drop in intronic GC content associated to most TEI and many NAHR deletions would increase the difference of GC content between introns and their flanking exons, what could facilitate exon definition during splicing and might contribute to the observed differential expression of some transcripts. It has been recently shown that human enhancers are associated to high GC [38] and that Alu elements can act as enhancers [39], suggesting that deletions could not only alter splicing but also influence regulatory features located within introns.

### Deletion of intronic regions and changes in expression in trans

Introns in human are particularly enriched in regulatory regions and frequently interact with gene promoters of other genes via chromatin looping (**Fig 4D**). Therefore, deletions in introns that show long-range interactions with promoters of other genes could potentially affect their expression (*trans* effects). We used promoter-capture Hi-C published data for B-lymphocytes [40] to link intronic regions and gene promoters. We identified 322 deletions in intronic regions that interact with gene promoters of other genes (672 in total). Taking all combinations of genes and the trans-intronic regions with deletions, we searched for intronic trans-DEL-eQTLs: intronic regions that, when deleted, are associated with changes in expression of a different gene. Twelve of these genes were found to be significantly differentially expressed in the individuals presenting an intronic deletion in another gene (trans-intron-eGenes, **Fig 4F-G**). For example, PRSS36 (Protease, Serine 36) is downregulated in individuals with a intronic deletion in SETD1A (SET Domain Containing 1A) gene, while LIAS (Lipoic Acid Synthetase) gene is upregulated in individuals with a intronic deletion in PDS5A (PDS5 Cohesin Associated Factor A) (**Fig 4G)**. In addition, 81 transcripts from 65 genes were also differentially expressed (trans-intron-eTranscripts) in the individuals with a trans-DEL-eQTLs. The loss of intergenic fragments in 3D contact with a gene were associated to a similar number of DEGs than the DEGs associated to intronic trans-DEL-eQTLs (16 trans-eGenes, 123 eTranscripts associated to intergenic deletions, **Fig 4F** and **Table S5**).

We analysed the age of different types of eGenes and observed that whole-gene and exonic eGenes are enriched in young age classes (**Fig 5A**). This pattern is very different in intronic and intergenic eGenes: intronic cis-eGenes are enriched in old ages, while intronic trans-eGenes and intergenic-eGenes do not seem to be associated with gene age. If we compare the RVIS of the different types of eGenes, we find that whole gene and exonic eGenes are actually among the most tolerant genes to point mutations in their coding sequence (**Fig 5B**). In contrast, we found that a significant proportion of intronic cis-eGenes with low RVIS percentiles, indicating that protein-coding genes that are intolerant to point mutations at the protein level can have intronic deletions associated to gene expression changes. Strikingly, trans-eGenes show the lowest RVIS percentiles, indicating that intronic variation might impact the gene expression of interacting genes that are quite intolerant to coding mutations (**Fig 5B**).

**Fig 5.**
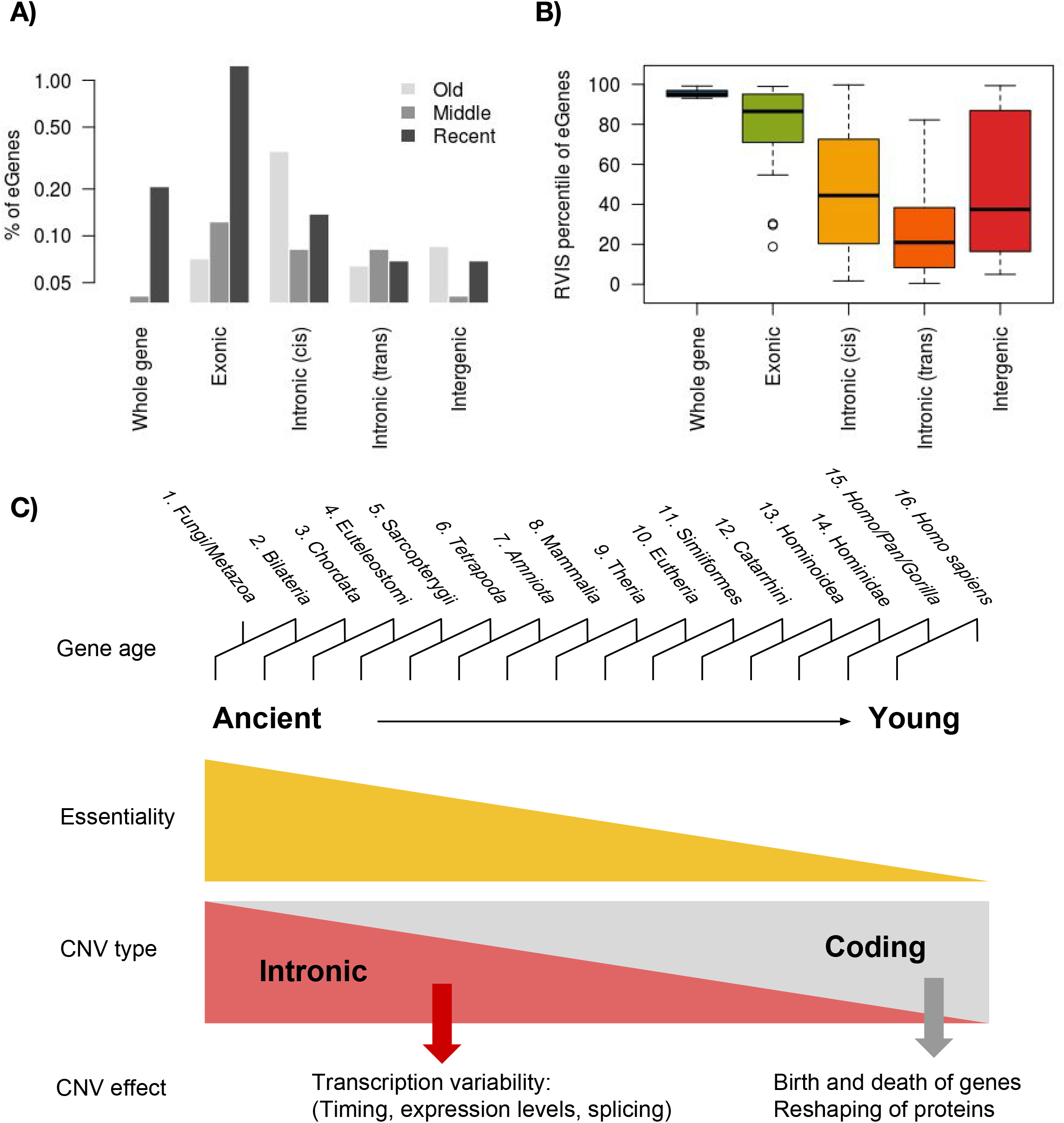
Impact of CNVs on genes and their evolution. (A) Percentage of genes of each group of evolutionary ages that is associated to an eCNV, for each type of eCNV. (B) RVIS percentile of the eGenes, by type of eCNV. Genes with the lowest percentile are among the most intolerant of human genes. (C) Evolutionarily ancient and young genes accumulate different kinds of structural variants. While young genes are enriched in coding deletions (which alter gene dosage or disrupt the protein, sometimes affecting gene expression), ancient genes have highly conserved coding sequence but an enrichment of deletions within their introns. As we have shown, these changes in introns can be associated with changes in gene expression, showing that although the protein is highly conserved, the expression of it can change from an individual to another due to changes in regulation.

### Intronic deletions can alter enhancer sequences or their location

To further study the potential impact of intronic deletions in regulatory regions, we analyzed the co-occurrence of these events with enhancers. In eGenes or eTranscripts, 15 intronic DEL-eQTLs overlap with enhancers (an overlap that is higher than expected by chance, *P* = 0.023, odds ratio = 2.04, Fisher’s test). These 15 deletions represent the 24% of the tested intronic deletions overlapping with enhancers in this cell type. We need to consider that many intronic deletions that were not investigated because they fall within genes that are expressed in other cell types. Based on our observations in lymphoblastoid cells, we estimate that there might be 105 additional eDels of the 422 that overlap with enhancers. Regarding the deletions not overlapping with enhancers, we found that the distance between the DEL-eQTL and the closest enhancer is shorter than the distance of the deletions not associated with expression changes (*P* = 9.2e-04, Student’s t-test). These results suggest that intronic DEL-eQTLs could also be affecting interactions between promoter and intronic enhancers without directly disrupting the enhancer sequence.

Motivated by these findings, we investigated if there is a global tendency (independently of gene expression) for intronic deletions to affect or not affect enhancers. First, we observed that enhancers are enriched in introns (P < 1e-04, global permutation test) agreeing with previous findings in plants [41,42]. Strikingly, we find that intronic deletions and intronic enhancers co-occur in the same intron more often than expected by chance (P < 2.2e-16, Fisher’s test), possibly because most intronic deletions and intronic regulatory features are found in very long introns. However, by randomly relocating each intronic deletion within the same intron, we observed that the direct overlap of the deletions with enhancers is significantly lower than expected (P = 0.0304, global permutation test,, **Table S6**). A possible functional interpretation of these results is that it might be some degree on plasticity on the distance between intronic enhancer and promoters, but many intronic enhancers might be essential and cannot be lost. Interestingly, as we saw above, the loss of non-essential intronic enhancers can be associated to changes of gene expression.

## Discussion

Intronic CNVs constitute the most abundant form of CNV in protein-coding genes (**Fig 1**) and might have a previously unsuspected role in human evolution and disease. This variation in intronic length in healthy human populations implies that the actual size of many genes is different among individuals and, therefore, it might change in populations over time. However, little attention has been given to this variability even if gene length has been shown to be important in many genes.

We have shown that that intronic deletions are occurring more often than expected by chance in three different CNV maps (using two different background models). Other studies have previously reported that CNVs are impoverished [20,43] or neither impoverished nor overrepresented within introns [44]. To explain this apparent controversy, we have to carefully review the different definitions of “intronic CNVs”. Here, we looked at deleted regions located completely within constitutive intronic regions (excluding intronic regions that contain alternative exons). Mu et al. [44] showed that purely intronic CNVs in general are not either enriched or impoverished in their dataset, but they observed that the subset of events associated to NAHR are found more often than expected by chance using the Pilot Phase dataset of the 1KG Project. We obtained similar results using the CNV map by Abyzov et al. [17] (Phase 1 of the 1KG) where intronic deletions were neither significantly enriched nor impoverished but the subset of NAHR deletions was significantly enriched (data not shown). These results illustrate the importance of a clear definition of intronic CNVs and the danger of generalising the results of one particular study. Each study is normally biased to detect different mechanisms, different sizes or types of CNVs or events observed at different frequency in distinct populations.

Our results suggest that copy number variation is shaping gene evolution in different ways depending on the age of genes, duplicating or deleting young genes and contributing to fine-tuning the regulation in both young and old genes (**Fig 5C**). Although we expect stronger functional effects for CNVs affecting the coding sequence, we have shown that the purely intronic CNVs also show signatures of positive selection. Indeed, the proportion of intronic CNVs with adaptive potential is similar or higher than for coding CNVs. Popadin et al. showed that primate-specific genes in human are enriched in single nucleotide variants correlated with gene expression (cis-eQTLs) with their associated SNPs tending to be closer to the TSS than in older genes [45]. These data highlight the need of dissecting the different types of genetic variation in order to understand the complex relationships between SNPs, CNVs, gene expression and gene age. While point mutations near the TSS [45] and coding CNVs seem to have a higher effect in young genes, intronic CNVs are frequently associated with gene expression variation in genes of any age.

Previously published studies on the effect of genetic variants on gene expression have proven the effect of CNVs on expression variability [46–48]. Chiang and co-workers identified 789 SVs associated to changes in gene expression, most of them (88.3%) not overlapping with exons from the eGene [48]. DeBoever and co-workers observed that a large proportion of common CNVs associated with gene expression levels is located in intergenic regulatory regions [47]. However, research on the subject has been mostly restricted to SVs found within 1Mb from the gene and previous works did not analysed intronic regions in detail. In contrast, we relied on Hi-C data to define deletions affecting regions in 3D contact with a gene. In this way, we do not require the CNV to be located within any particular distance to the TSS position of a gene. We tested intergenic eCNVs that can be located at any distance from 864bp to 82Mb from the nearest gene.

The differences that we observe in gene expression could be the result of intronic CNVs affecting the rate of transcription, the splicing process, the stability of RNA or a combination of them. For example, intronic deletions interfering with splicing recognition might trigger the nonsense-mediated decay (NMD) pathway that would degrade the transcript. Recently, it has been shown that the balance of unspliced and spliced mRNA (RNA velocity) is a cell type-specific signature that can be used to predict the future increases or decreases of gene expression in single cells [49]. As the amount of unspliced transcript detected will depend on the length of introns - which can be highly variable in some genes - we would expect that the RNA velocity of intron-varying genes will be also varying in human populations.

Despite the clear trends shown, our results are likely to underestimate the extent of the impact of intron loses in gene expression. On one hand, we only investigated the effect on gene expression in lymphoblastoid cell lines. On the other hand, the regulatory data currently available is also limited. The interaction maps change in different cell types [40,50] and many enhancers are tissue-specific [51]. Therefore, the loss of intronic sequence could affect the expression of genes in other cell types. In addition, the 3D contacts involving frequently deleted regions in the population will be underrepresented in the interaction map used in our study, as they are less likely to be present in the assayed samples. The availability of CNV, personal gene expression and genome interactomes from multiple tissues will allow to evaluate more accurately what is the impact of coding and non-coding deletions in the whole organism.

### Supplemental Data

Supplemental Data include twelve figures and eight tables, including a list of all CNVs used in this study and a list of genes overlapping these CNVs.

## Methods

### Origin and filtering of CNV maps

Whole genome CNV maps were downloaded from 5 different publications [17–21]. For our analysis we selected autosomal and not private CNVs. Some extra filters were applied to some maps: In Handsaker et al. we removed CNVs marked as low quality and all the variants from two of the individuals (NA07346 and NA11918) because they were not included in the phased map. From Zarrei’s maps we used the stringent map that considered CNVs that appeared in at least 2 individuals and in 2 studies. The complete list of CNVs analysed is available in **Table S7.**

### Gene structures

Autosomal gene structures and sequences were retrieved from Ensembl [52] (http://www.ensembl.org; version 75) and principal isoforms were determined according to the APPRIS database [53], Ensembl version 74. In order to avoid duplicate identification of introns, intronic regions were defined as regions within introns that aren’t coding in any transcript of any gene. When analyzing real introns, in order to avoid duplicate identification of introns, the principal isoform with a higher exonic content was taken. The complete list of genes affected by different types of CNVs is available in **Table S8**. Genomic sequences were obtained from the primary GRCh37/hg19 assembly, and were used for calculating the GC content of introns and intronic CNVs.

### Essential genes

The list of essential genes was obtained by aggregating lists of genes reported as essential after CRISPR-based genomic targeting [54,55], gene-trap insertional mutagenesis methodology [56], and shRNA [57–59].

### Dating gene and intron ages

An age was assigned to all duplicated genes as described before [27]. In the case of singletons gene ages were assigned from the last common ancestor to all the genes in their family according to the gene trees retrieved from ENSEMBL. Singleton’s ages can be noisy for genes suffering important alterations as gene fusion/fission events or divergence shifts. As a consequence, these ages should not be interpreted as the age of the oldest region of the gene, but as a restrictive definition of gene age considering a similar gene structure and gene product.

The ages (from ancient to recent) and number of genes per age are as follows: FungiMetazoa: 1119, Bilateria: 2892, Chordata: 1152, Euteleostomi: 8230, Sarcopterygii: 182, Tetrapoda: 154, Amniota: 408, Mammalia: 375, Theria: 515, Eutheria: 848, Simiiformes: 233, Catarrhini: 170, Hominoidea: 106, Hominidae: 64, HomoPanGorilla: 204, HomoSapiens: 500. For some analyses, Primates age groups (Simiiformes to HomoSapiens) were collapsed. For other analyses, we only grouped the 16 ages in three, “ancient” (collapsing groups from FungiMetazoa to Sarcopterygii), “middle” (from Tetrapoda to Eutheria) and “young” genes (Primates).

Intronic regions were assigned the evolutionary age of the gene they belonged to. In the cases when an intron could be assigned to more than one gene, the most recent age was assigned to them.

### Statistical assessment of genome-wide distribution of CNVs

To estimate statistical significance of our results we performed permutation tests. In order to compare the number of overlaps of CNVs with genic functional elements we compared our observed values to a background model. A global background was obtained by relocating all the CNVs in the whole genome 10,000 times, avoiding low-mappability regions in R package “BSgenome.Hsapiens.UCSC.hg19.masked”). Genome coordinates and low mappability regions were downloaded using RegioneR package [60]. A local background was obtained by segmenting the genome in 278 windows of at least 10Mb and randomly shuffling the CNVs within their original window 10,000 times, also avoiding low-mappability regions. P-values were computed using a function derived from the permTest function from package RegioneR version 1.6.2 [60]. Code is available in https://github.com/orgs/IntronicCNVs.

We compared the location of the CNVs in our datasets and compared with their distribution in the random models in order to calculate enrichments or depletions depending on the intron size and gene age and essentiality.

### Regulatory features

We downloaded a genome-wide set of regions that are likely to be involved in gene regulation from the Ensembl Regulatory Build [61], assembled from IHEC epigenomic data [62]. We checked if introns are enriched in these regulatory features (promoters, enhancers, promoter flanking regions or insulators) by comparing to a random background model generated by relocating 10,000 times all regulatory features in the genome. P-values are the fraction of random values superior or inferior to the observed values.

In order to check for the significance of the overlaps between intronic deletions and regulatory features we relocated 10,000 times each intronic deletion within their host intronic region, avoiding overlaps with exons. Then, we compared the observed and the expected overlap with regulatory features. Introns that overlapped with low-mappability regions were previously removed.

### Alu elements

Alu element genomic coordinates were extracted from the RepeatMasker tracks from UCSC, build GRCh37.

### CNV mechanisms

The analysis intronic deletions generated through different mechanisms was done using the dataset from Abyzov’s [20] study.

### Gene expression analysis

We used available RNA-seq data at Geuvadis [33] that was derived from lymphoblastoid cell lines for 445 individuals who were sequenced by the 1KG Project and for whom we have the intronic deletions in the largest CNV map [20]. We focused our analyses on the 763 genes that have only one intronic deletion in the population with at least two individuals affected in the Geuvadis dataset. For each of these genes we classified the PEER normalized gene expression levels [63] in two groups: 1) gene expression of individuals homozygous for the reference genotype and 2) gene expression of individuals with one allele with the deletion and the other with the reference genotype. We then performed Student’s t-tests to compare the expression of the two different genotypes. We corrected for multiple testing with p.adjust R function (Benjamini-Hochberg method). In addition, in order to see if the number of significant differentially expressed genes is higher than expected by chance, for each intronic deletion, we the shuffled 10,000 times the genotypes of the individuals and performed t-tests with the expression of the random groups of wild-type and heterozygous individuals. For example, if a deletion is found in heterozygosis in 50 individuals and the rest are wild-type, we will test if there is differential expression when comparing the expression of 50 randomly selected individuals versus the rest. By repeating this shuffling 10,000 times for every tested deletion we can calculate the expected percentages of significantly differentially expressed genes.”

### Observed vs expected intronic deletion content score

The number and size of expected intronic deletions per gene was calculated in two different ways: 1) relocating 10,000 times all deletions in the whole genome (except for low mappability regions) and 2) relocating 1,000 all intronic deletions within the intronic regions. In both cases, a score was generated to determine what genes have more or less intronic deletions than expected. This score was calculated taking into account 1) the ranked position of the number of intronic deletions per gene divided by their median expected value, 2) the ranked position of the observed divided by the median expected size of the deletions, 3) the ranked position of the percentage of intronic content that is lost, 4) the ranked inverse of the expected intronic loss and 5) the ranked frequency of the deletion in the 1KG Project populations, if available. Because the frequency of the event depends on the reference genome, we find that a deletion present in, for example, all except for two individuals, should probably be considered as a rare gain and the deletion should be the reference. For this reason, the values were normalized in a way that 0.5 would be the maximum frequency and 0.9 and 0.1 would be given the same position in the ranking. Once all rankings were calculated and normalized from 0 to 1, a score was assigned to each gene by averaging their five ranks. The final set of 458 genes with less deletions than expected is the intersection of the top 500 genes of the two randomizations, and the set of 484 genes with more deletions than expected, the intersection of the bottom 500 genes.

### Functional enrichment analysis

Functional enrichment analysis of the genes with a lower scores and higher scores was performed with DAVID [31] and STRING [32]. Enrichment of essential genes in our datasets was performed with a Fisher test using our list of essential genes (see the “Essential genes” section in Materials and Methods).

### Population stratification

For the study of population stratification of deletions, Vst statistics were extracted from Sudmant Nature [20]. As in Sudmant Nature, a cutoff of 0.2 was selected to indicate high population stratification of a locus.

## Acknowledgments

The authors thank Salvador Capella, Matthew Bashton, Venetia Bigley, Aneta Mikulasova, Joris Veltman, Tomas Marques-Bonet, Vera Pancaldi, Inmaculada Hernandez and Ruth Rodríguez-Barrueco for their constructive comments on the manuscript. Alfonso Valencia acknowledges the Joint BSC-CRG-IRB Research Program in Computational Biology and Daniel Rico thanks the Newcastle University NURF programme and the Centre for Health and Bioinformatics. Maria Rigau acknowledges a La Caixa fellowship.

This work was supported by Project Retos BFU2015-71241-R Spanish Ministry of Economy, Industry and Competitiveness (MEIC, http://www.mineco.gob.es), co-funded by European Regional Development Fund (ERDF) to A.V. and Wellcome Trust (https://wellcome.ac.uk) Seed Award in Science (206103/Z/17/Z) to D.R.

## Supplementary figures and tables legends

**Figure S1. Comparison of datasets**

Only variants in autosomes are considered and private events are excluded. (A) Number and type of CNVs per dataset. (B) Autosomal Mb that are CNV. Gray part of the bars corresponds to the CNV Mbs that are shared among maps. Colored parts of the bars are map-specific CNV regions. (C) Width distribution of gains and losses in each map bean lines and overall line are means). (D) Number of subjects and number of populations of origin used for building of each filtered map. DGV: Database of Genomic Variants (http://dgv.tcag.ca). For more information on the 1000 Genomes Project, see http://www.internationalgenome.org.

**Figure S2. Enrichment analysis of purely intronic, intron-intersecting and coding deletions**

Enrichment or impoverishment of deletions within introns, deletions intersecting introns (purely intronic and intron-exon combined) and exon-overlapping deletions (purely coding and intron-exon combined) in different maps of copy number variation and using global (A) and local (B) background models. Values are given as log2 ratios observed/expected (median expected value from 10,000 randomisations). Error bars show the median absolute deviation and asterisks indicate significance: * for P<0.05, ** for P<0.005 and *** for P<0.0005.

**Figure S3. Enrichment analysis of deletions in introns of different sizes**

Ratios of observed versus expected number of deletions in each size bin after 10,000 random permutations using global (A) and local (B) background models. All size bins have a similar number of intronic regions (deciles, size intervals indicated between brackets). Asterisks mark the bins significantly enriched with intronic deletions: * for P<0.05, ** for P<0.005 and *** for P<0.0005.

**Figure S4. Enrichment analysis of CNVs (including gains, losses and gain/losses) in genes of different evolutionary ages**

Ratios of observed versus expected number of genes with CNVs (gains, losses and gain and loss CNVs) affecting their coding region in each gene age after 10,000 random permutations using the global (A) and local (B) background models. Abyzov et al. (2015) is excluded because it is the only CNV map that do not contain any gain. Red asterisks show an enrichment when above the box, a depletion when below the box: * for P<0.05, ** for P<0.005 and *** for P<0.0005.

**Figure S5. Impact of deletions on genes of different evolutionary ages.**

Percentage of genes from each gene evolutionary age that contain intronic deletions in (A) or deletions overlapping with exons (including partial and whole gene CNVs) in (B). The gray line represents the expected value, calculated as the median of the genes in the 10,000 random permutations. Significance is marked with asterisks: * for P<0.05. ** for P<0.005.*** for P<0.0005 and their color represents enrichment (red) or impoverishment (black).

**Figure S6. Differential effect of intronic deletions in big introns**

Ratios of observed versus expected number of deletions within in introns bigger than 1.5kb from different evolutionary ages. Expected values are calculated 10,000 random permutations using a global background model. Asterisks show an enrichment when above the box, a depletion when below the box: * for P<0.05, ** for P<0.005 and *** for P<0.0005.

**Figure S7. Effect of the different types of deletions on all evolutionary ages.**

Proportion of genes with deletions that have the whole locus deleted, only part of their exons (exonic) affected by deletions or intronic deletions only. This figure is equivalent to Figure 3C, but here separated by CNV map.

**Figure S8. Essential genes**

(A) Intron sizes of non-essential and essential genes. (B) Percentage of essential genes per evolutionary age.

**Figure S9. Highly stratified deletions associated with expression differences**

(A) Characteristics of highly stratified variants (HSVs) that are significant cis-intronic-eDeletions. (B) Gene expression of “wild-type” (CN=2) individuals and heterozygous carriers (CN=1) of the eDeletion.

**Figure S10. GC content in introns and intronic deletions**

(A) Bean-plots showing the different GC distribution between the flanking exons of introns with or without deletions, separated by intron size bins (with equal number of introns per bin). (B) GC content distributions in introns with or without deletions, separated by intron size bins. Significance is considered for p-values < 0.05. Beans show the estimated density of each distribution; horizontal lines show the mean values of each side of the bean and the dashed horizontal line line represents the overall average of all values.

**Figure S11. Examples of introns with a drop of GC content**

X-axis represents the coordinates of the intron with its flanking exons (black boxes). Y-axis shows the GC content, calculated with sliding 200 bp windows. The deleted region is highlighted in grey.

**Figure S12. Alu element content in regions deleted by different mechanisms**

(A) Proportion of deletions of each mechanism that overlaps with Alu elements. (B) Percentage of the deleted regions covered by Alu elements. Deleted regions and mechanisms from Abyzov et al. (2015). NAHR: Non-allelic homologous recombination. NH: Non-homologous end joining. TEI: Transposable Element Insertion.

**Table S1. Number of individuals in each map, project the variants belong to and methods used for CNV detection.**

**Table S2. Fold changes and P-values for all maps**

**Table S3. Top genes with more or less deletions than expected**

**Table S4. List of differentially expressed genes (eGenes)**

**Table S5. Summary of differentially expressed transcripts (eTranscripts)**

**Table S6. Overlap of intronic deletions with regulatory features.**

**Table S7. List of CNVs and their impact on genes**

**Table S8. Genes affected by deletions in any CNV map**

